# Receptive field formation by interacting excitatory and inhibitory synaptic plasticity

**DOI:** 10.1101/066589

**Authors:** Claudia Clopath, Tim P. Vogels, Robert C. Froemke, Henning Sprekeler

**Affiliations:** Bioengineering Department, Imperial College London, South Kensington Campus London SW7 2AZ, UK; Centre for Neural Circuits and Behaviour, University of Oxford, Mansfield Road Oxford OX1 3SR, UK; New York University, School of Medicine, 540 First Avenue New York, NY 10016, USA; Dep. for Electrical Engineering and Computer Science, Berlin Institute of Technology and Bernstein Center for Computational Neuroscience Marchstr. 23, 10587 Berlin, Germany

## Abstract

The stimulus selectivity of synaptic currents in cortical neurons often shows a co-tuning of excitation and inhibition, but the mechanisms that underlie the emergence and plasticity of this co-tuning are not fully understood. Using a computational model, we show that an interaction of excitatory and inhibitory synaptic plasticity reproduces both the developmental and – when combined with a disinhibitory gate – the adult plasticity of excitatory and inhibitory receptive fields in auditory cortex. The co-tuning arises from inhibitory plasticity that balances excitation and inhibition, while excitatory stimulus selectivity can result from two different mechanisms. Inhibitory inputs with a broad stimulus tuning introduce a sliding threshold as in Bienenstock-Cooper-Munro rules, introducing an excitatory stimulus selectivity at the cost of a broader inhibitory receptive field. Alternatively, input asymmetries can be amplified by synaptic competition. The latter leaves any receptive field plasticity transient, a prediction we verify in recordings in auditory cortex.

## Introduction

The balance of excitatory and inhibitory currents (E/I balance) received by cortical neurons (Wehr & Zador, 2003; Monier et al., 2008) is thought to be essential for the stability of cortical network dynamics and provides an explanation for the irregular spiking activity observed in vivo (van Vreeswijk & Sompolinsky, 1996; Renart et al., 2010; Ecker et al., 2010). Although the balanced state is a relatively robust dynamical regime in recurrent networks with random connectivity (van Vreeswijk & Sompolinsky, 1996), the mechanisms by which it is maintained in the presence of synaptic plasticity on virtually all synaptic connections in the mammalian brain (Malenka & Bear, 2004) are poorly understood. Activity-dependent Hebbian plasticity of inhibitory synapses has been suggested as a self-organization principle by which inhibitory currents can be adjusted to balance their excitatory counterparts (Vogels et al., 2011; Luz & Shamir, 2012).

E/I balance also shapes responses of single cells to sensory stimulation (de la Rocha et al., 2008; Carvalho & Buonomano, 2009; Froemke et al., 2007; Vogels et al., 2011). This control of neuronal output by the interplay of excitation and inhibition suggests that through the establishment of E/I balance inhibitory synaptic plasticity can in turn exert an influence on excitatory plasticity (Wang & Maffei, 2014).

Excitatory synaptic plasticity is thought to form the basis of receptive field development in sensory cortices. Stimulus-specific receptive fields require a spontaneous symmetry breaking, i.e., an impromptu departure from equally weighted inputs in favor of a few strong ones. Such symmetry breaking can be achieved by competitive interactions, either between neurons (Kohonen, 1982; von der Malsburg, 1973) or between the afferent synapses onto a given neuron. Synaptic competition can be realized through synaptic learning rules of a simple Hebbian (Linsker, 1986; Miller, 1995; Wimbauer et al., 1997) or Bienenstock-Cooper-Munro (BCM) type (Bienen-stock et al., 1982; Clopath et al., 2010). In the former, synaptic competition is often amplified by an additional weight-limiting mechanism. In contrast, BCM rules rely on a sliding threshold between potentiation and depression that depends on the recent activity of the neuron.

These theories for receptive field formation have mostly considered purely excitatory networks or do not respect Dale’s law, and thus cannot reproduce the correlated stimulus tuning (co-tuning) of excitatory and inhibitory currents that is observed in sensory cortices (Wehr & Zador, 2003; Froemke et al., 2007; Anderson et al., 2000; Monier et al., 2008; Harris & Mrsic-Flogel, 2013). Here we investigate under which conditions neurons can develop stimulus selectivity and E/I co-tuning simultaneously (Wehr & Zador, 2003; Froemke et al., 2007). To this end, we analyze the dynamical interaction of excitatory and inhibitory Hebbian plasticity. A combination of mathematical analysis and computer simulations shows that the determining factors controlling the dynamics of receptive field development are i) the time scales of excitatory and inhibitory plasticity, ii) the stimulus tuning width of the inhibitory inputs and their excitatory counterparts and iii) possible activity biases in the input.

We show that plastic inhibitory inputs with a broad stimulus tuning can functionally serve as a sliding threshold and generate BCM-like behavior. In contrast, narrowly tuned inhibitory inputs lead to a detailed balance (Vogels & Abbott, 2009) of excitatory and inhibitory currents and thereby exert an equalizing effect on the postsynaptic neuronal response rather than the competition induced by the sliding threshold. In this case, the simultaneous establishment of a receptive field and E/I co-tuning can be induced by small heterogeneities in the inputs. By combining the interaction of excitatory and inhibitory plasticity with neuromodulation-induced dis-inhibition (Froemke et al., 2007; Dorrn et al., 2010; Letzkus et al., 2011), our model reproduces a wide range of dynamical phenomena that arise during receptive field plasticity in the auditory cortex (Froemke et al., 2007).

## Results

We study the interaction of excitatory and inhibitory synaptic plasticity in a single postsynaptic rate-based model neuron receiving both excitatory and inhibitory synaptic inputs. Unless otherwise mentioned, the neuron receives a set of sensory stimuli, each of which activates a separate presynaptic excitatory neural population. Excitatory synapses are plastic according to a Hebbian rule, i.e., the change of the synaptic weight 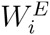 from excitatory input population *i* is proportional to the product of presynaptic population activity *E_i_* and postsynaptic activity *R*. Because Hebbian plasticity in excitatory synapses is unstable, this rule is combined with a weight-limiting mechanism, in the form of a subtractive or multiplicative weight normalization:

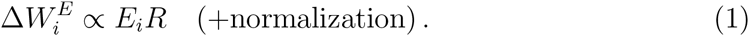

Subtractive normalization reduces all excitatory weights by the same amount, such that the sum of the weights is held constant, phenomenologically mimicking a competition of all excitatory synapses for a fixed pool of postsynaptic receptors. Multiplicative normalization scales all weights such that the sum of the squared weights is held constant, a mechanism that could be implemented by homeostatic synaptic scaling (Turrigiano et al., 1998). As known from previous studies, the choice of the normalization can have a strong impact on the learning dynamics, because it determines the degree of competition between different excitatory synapses (Miller & MacKay, 1994; Dayan & Abbott, 2001).

The inhibitory synapses onto the neuron are subject to a Hebbian plasticity rule that changes their synaptic weight in proportion to presynaptic activity and the difference between postsynaptic activity and a target rate *ρ*_0_. This rule has previously been shown to balance excitation and inhibition such that the firing rate of the postsynaptic cell approaches the target firing rate *ρ*_0_ (Vogels et al., 2011).

Here, we study the emergence of excitatory and inhibitory receptive fields in sensory cortices, as a result of the interaction of excitatory and inhibitory synaptic plasticity. We investigate the learning dynamics of this model for different sensory input profiles and relative learning rates of excitatory and inhibitory plasticity.

### Unspecific inhibition sets a sliding threshold and mediates BCM-like receptive field formation

We first consider a situation in which the inhibitory afferents have no stimulus tuning, but rather constant firing rates (Figure 1A). Inhibitory synaptic plasticity is assumed to act more rapidly than excitatory plasticity. Under these conditions, inhibitory plasticity establishes a state of a *global* E/I balance (Vogels & Abbott, 2009; Vogels et al., 2011), in which inhibition balances excitation on *average* across stimuli. Because inhibitory plasticity is rapid, this balance is dynamically maintained in the presence of excitatory changes (Vogels et al., 2011).

**Figure 1:**
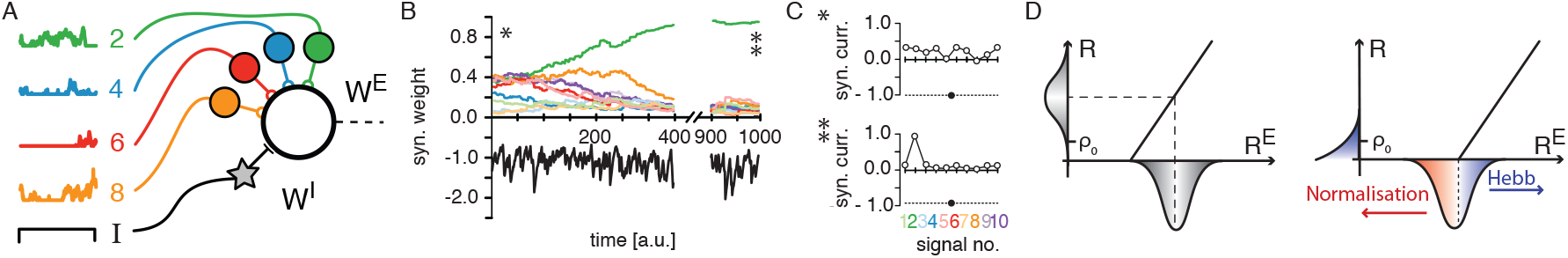
Receptive field formation with unspecific inhibition. A) Network diagram. A single postsynaptic neuron receives synaptic input from 10 excitatory populations (colored circles, not all 10 shown) with different time-varying firing rates (signals, colored traces) and from a single inhibitory population (grey star) with constant firing rate (black trace). B) Temporal evolution of excitatory (positive) and inhibitory (negative) synaptic weights. C) Synaptic currents evoked by activating individual signals before (top) and after learning (bottom). Stars indicate corresponding times in B. D) Illustration of the mechanism that leads to the emergence of selectivity in the synaptic weights (D) before E) after convergence). Results for multiplicative normalization, for subtractive normalization see Figure 1

For such unspecific inhibition, the interaction of excitatory and inhibitory plasticity leads to a robust development of a receptive field with high stimulus selectivity (Figure 1B, C). The underlying mechanism is similar to that of BCM learning rules (Bienenstock et al., 1982). In BCM rules, a sliding threshold in postsynaptic activity dictates the sign of plasticity, causing a competition between different stimuli and providing a homeostatic mechanism for the postsynaptic firing rate. In our case, inhibitory plasticity acts as a similar homeostatic mechanism that rapidly adapts the strength of the inhibitory input such that the mean firing rate of the postsynaptic cell is near the target rate *ρ*_0_ (Figure 1, right). For strong excitatory inputs (Figure 1D, histogram below horizontal axis), inhibition will dominate for most stimuli. Only a few stimuli that activate sufficiently strong excitatory synapses can evoke postsynaptic activity (Figure 1D, right). Only those synapses will experience coincident pre- and postsynaptic activity and will be potentiated by the Hebbian learning rule (Figure 1D, blue arrow). All others will be suppressed by the normalization (Figure 1D, red arrow). Consequently, stimuli that evoke a strong response are rewarded and all others are punished, resulting in the formation of a strong stimulus selectivity (Figure 1C).

A mathematical analysis of the learning dynamics makes this intuition explicit. Assuming a linear output neuron with rectification, the cell can only be active when the total excitatory input 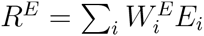 exceeds the total inhibitory input *θ* = *W^I^ I* (where *W^I^* is the total inhibitory synaptic strength and *I* the constant activity of the inhibitory input). Because the Hebbian learning rule Eq. 1 for the excitatory weights requires postsynaptic activity, excitatory plasticity is effectively thresholded by the activity-limiting inhibitory input and can be written as:

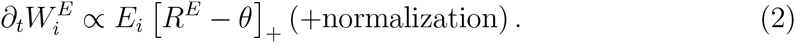

The threshold *θ* is sliding, because the inhibitory plasticity perpetually adjusts the inhibitory weights such that the mean output rate of the cell is equal to the target rate *ρ*_0_ (see SOM for a mathematical derivation).

The effective learning rule Eq. 2 has the form of a BCM rule in that synaptic changes are proportional to the product of presynaptic activity and a nonlinear function of the total excitatory drive. There are also differences to BCM theory. In particular, BCM rules induce synaptic depression below the threshold, while the effective learning rule Eq. 2 has no explicit depression component (Figure 1D). Instead, depression is a side-effect of the synaptic weight normalization. Nevertheless, the mechanism by which the system establishes the receptive field is the same as in BCM rules: a sliding threshold that introduces a temporal competition between stimuli. This mechanism does not rely on the assumption of constant inhibitory input rate, it also applies when the inhibitory input is pooling over all excitatory inputs (Figure 1 - Supplement 1). Furthermore, the normalization procedure does not alter these results.

### Egalitarian effects of stimulus-specific inhibition

Experimental evidence from different sensory systems indicates a stimulus co-tuning of excitatory and inhibitory currents (Wehr & Zador, 2003; Froemke et al., 2007; Anderson et al., 2000; Monier et al., 2008), which cannot be achieved with unspecific inhibition. We thus introduced stimulus-specific inhibitory inputs to investigate the formation of co-tuned receptive fields (Figure 2). We first studied a situation in which every excitatory input has an inhibitory counterpart with the same time-dependent firing rate (Figure 2A). Again, inhibitory plasticity is assumed to act more rapidly than excitatory plasticity.

**Figure 2:**
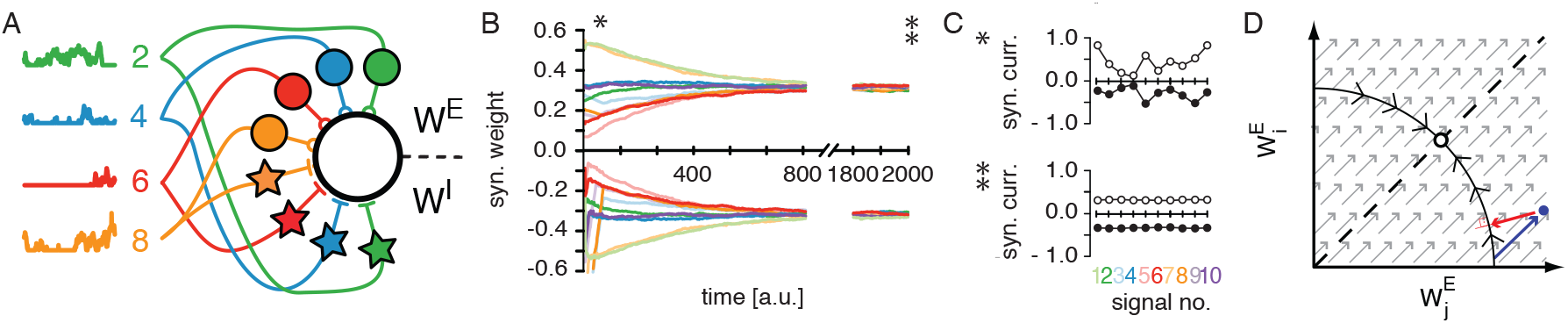
Receptive field formation with specific inhibition. A) Network diagram. A single postsynaptic neuron receives synaptic input from 10 excitatory populations (colored circles, not all 10 shown) and 10 inhibitory populations (colored stars). Each excitatory population has a different time-varying firing rate that is shared with a corresponding inhibitory population. B) Temporal evolution of excitatory (positive) and inhibitory (negative) synaptic weights. C) Synaptic weights before (top) and after learning (bottom). D) Illustration (for two excitatory weights only) of the mechanism that abolishes the selectivity in the synaptic weights. Because inhibitory plasticity equalizes the postsynaptic responses to all stimuli, the Hebbian excitatory rule increases all excitatory weights by the same amount (grey arrows, blue arrow: an example for such a Hebbian weight update). These changes are partly counteracted by the normalization that rescales the weight vector to unit length (grey arc, red arrow: example for normalization update), leading to an effective weight change that follows the black arrows along the constraint line. The joint dynamics drive all weights to the homogeneous fixed point (black circle).

For this highly stimulus-specific inhibition, all excitatory weights converge to the same value, and no receptive field emerges (Figure 2B, C). The inhibitory plasticity rule aims to establish a stimulus-specific detailed balance, with mean firing rates that are close to the target rate *ρ*_0_ for all stimuli individually (Vogels et al., 2011). For rapidly-acting inhibitory plasticity, this state is perpetually maintained in spite of (slower) synaptic changes in excitation. Mathematically, we can thus replace the postsynaptic firing rate in the excitatory learning rule by the target rate. This leads to an effective excitatory learning rule (see SOM for a more precise mathematical derivation):

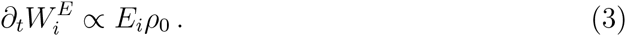

Because inhibitory plasticity forces the output of the neuron to the target rate *ρ*_0_, the dependence of the learning rule Eq. 1 on postsynaptic activity, and thus the Hebbian nature of the learning process is effectively lost and excitatory synaptic plasticity is driven by presynaptic activity only. If all input neurons fire at the same mean rate, all synapses undergo the same change on average. For a multiplicative normalization, which reduces large weights more than small weights (Figure 2D), this causes all excitatory synapses to converge to the same value. This is supported by a mathematical analysis (see SOM) showing that the homogeneous weight configuration is stable for a multiplicative normalization and changes only gradually when the firing rate of one individual input signal is increased (SOM, Figure 5 - Supplement 1). Thus, the interaction of excitatory and inhibitory plasticity on stimulus selective excitatory *and* inhibitory inputs does not favor the emergence of a receptive field. Interestingly, receptive field formation can nevertheless be reached in a subtractive normalization scheme, as shown below.

### Effects of the relative learning rates of excitation and inhibition

The effective learning rules Eqs. 2 and 3 were based on the assumption that the inhibitory plasticity is faster than its excitatory counterpart. As previously shown, this does not lead to the emergence of a receptive field when inhibition is stimulus-specific (Figure 3A-C). We wondered if a receptive field could emerge if the excitatory learning rate is increased, such that an excitatory stimulus selectivity is formed before inhibitory plasticity can establish a detailed balance and equilibrate the output to the target rate *ρ*_0_. Indeed, a stimulus-specificity emerges in the receptive field with increasing excitatory learning rate, but it is not stable. Instead, the weights start to show oscillations around the homogeneous state (Figure 3D-F), with an oscillation amplitude that increases with the excitatory learning rate (Figure 3D/F vs. G/I). This instability generates an intermittent turn-over of transient receptive fields (Figure 3G-I). In summary, with increasing excitatory learning rates, the synaptic weight configuration starts to show the emergence of transient receptive fields, at the cost of decreasing stability and precision of the detailed balance (Figure 3J).

**Figure 3:**
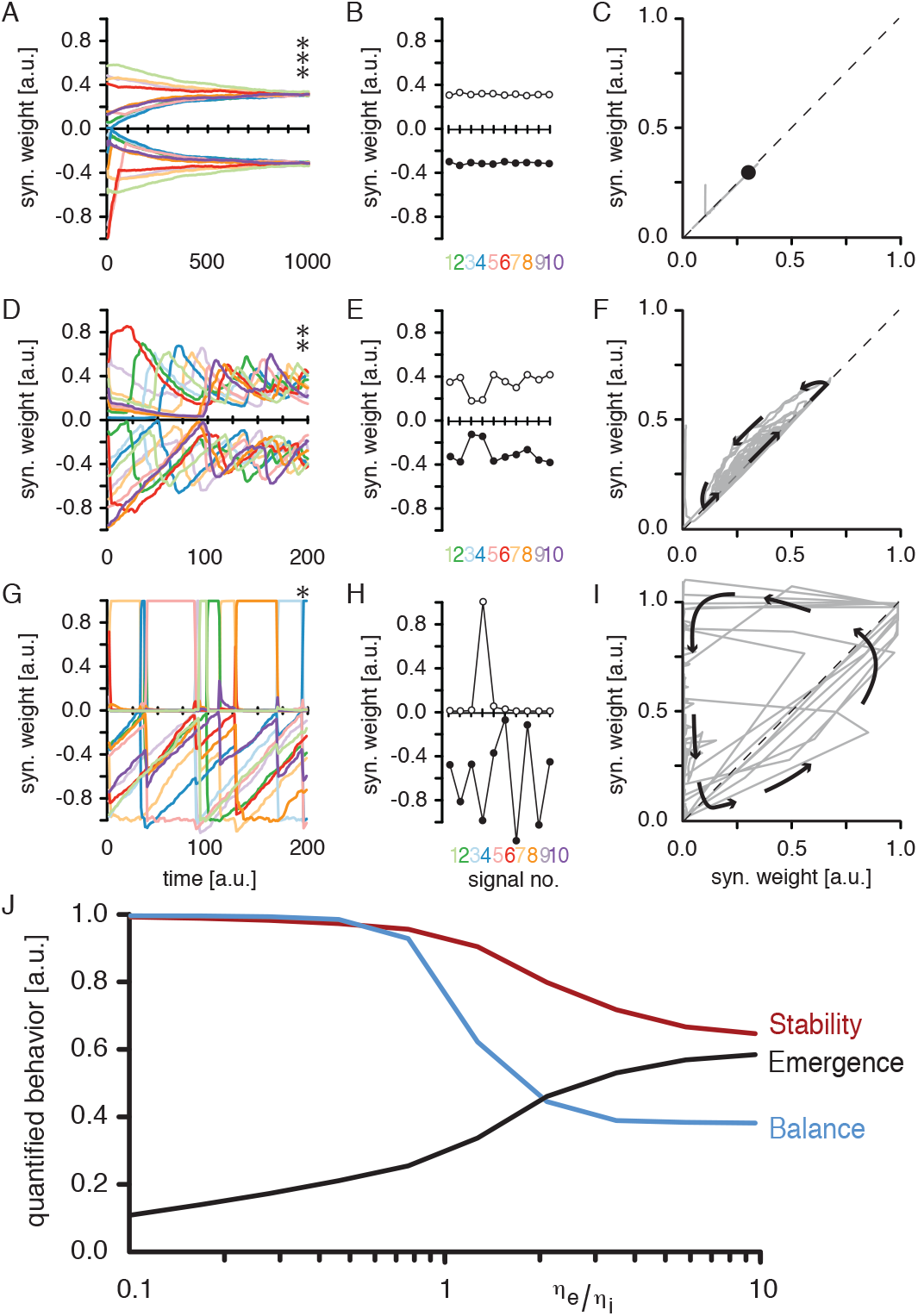
Effects of relative excitatory and inhibitory learning rate. A, D & G) Temporal evolution of excitatory (positive) and inhibitory (negative) synaptic weights, for low (A), intermediate (D) and high (G) ratio *η_E_*/*η_I_* of the excitatory inhibitory learning rates (*η_I_* is fixed and *η_E_* varies). B, E & H) Synaptic weights after learning (time indicated by star above A, E & G). C, F & I) Dynamics of the excitatory (horizontal axis) vs. inhibitory (vertical axis) synaptic weights for a selected input signal. For rapid inhibition, the inhibitory weights track their excitatory counterpart, all points are close to the diagonal. As the learning rate increases, increases in excitation trigger delayed increases in inhibition that restore the E/I balance and cause the excitatory weights to decay again. This causes a cyclic excursions in the excitatory-inhibitory weight plane, with increasing amplitude as the ratio *η_E_*/*η_I_* of excitatory and inhibitory learning rate increases. J) Dependence of the stability, balance and emergence indices of the weight configuration on the ratio *η_E_*/*η_I_* of excitatory and inhibitory learning rates.

The mechanism behind the observed oscillation is a delayed negative feedback loop on the stimulus selectivity that is introduced by the inhibitory plasticity. After the excitatory weights have converged to a selective state, inhibitory plasticity gradually establishes a detailed E/I balance and thereby “equilibrates” the neural responses to the different stimuli at the target firing rate *ρ*_0_. Once the postsynaptic response loses its stimulus selectivity, however, it can no longer support the existing receptive field and the neuron starts to fall back to the homogeneous weight configuration. In particular, the excitatory synaptic weights for the preferred stimulus start to decrease. Because the slow inhibitory plasticity lags behind, the associated inhibition remains strong, such that the previously preferred stimulus now becomes the least effective. As a result, a different stimulus is selected, albeit only until the inhibitory inputs for this stimulus have become sufficiently strong. These observations are supported by a linear stability analysis of the homogeneous, i.e., unselective weight configuration, which shows that the learning dynamics undergo a Hopf bifurcation as the ratio of the excitatory and inhibitory learning rates is increased beyond a critical value (see SOM).

### Broadened inhibitory tuning supports formation of receptive fields and broadened co-tuning

Unspecific inhibition supports receptive field emergence, but it can only generate a global E/I balance (Figure 1), which is inconsistent with the experimentally observed stimulus (co-)tuning of the inhibitory currents (Wehr & Zador, 2003; Froemke et al., 2007; Anderson et al., 2000). On the other hand, inhibitory plasticity of stimulus-specific inhibition, which does in principle allow a detailed E/I balance, generates a rate homeostasis for individual stimuli that hinders the emergence of a receptive field (Figure 2). Although the preferred stimuli for excitation and inhibition are similar in various sensory systems, the width of the inhibitory tuning varies substantially (Harris & Mrsic-Flogel, 2013). Hence, we hypothesized that inhibitory inputs with a stimulus tuning that is finite but broader than their excitatory counterparts – as is encountered, e.g., in visual cortex (Kerlin et al., 2010; Hofer et al., 2011) – could support both the formation of a receptive field and an (albeit degraded) detailed balance. To test this hypothesis, we assigned Gaussian input tuning curves to both the excitatory and inhibitory inputs, with tuning widths *σ_E_* and *σ_I_*, respectively (Figure 4). By controlling the inhibitory tuning width *σ_I_*, we can emulate the cases of highly specific inhibition (*σ_I_* ≤ *σ_E_*, Figure 4A-C), as well as cases of intermediate (*σ_I_* ≈ *σ_E_*, Figure 4D-F) and unspecific inhibition (*σ_I_* ≫ *σ_E_*, Figure 4G-I). Narrow inhibition (*σ_I_* = *σ_E_* = 1) allows a detailed balance, so that all weights converge to the same strength (Figure 4B, C), as expected from the earlier results. For very broad inhibition (*σ_I_* = 100), the excitatory weights show a spontaneous symmetry breaking, such that only few weights are large and most are zero (Figure 4H, I). For intermediate inhibitory tuning width, a receptive field emerges in the excitatory weights, and the inhibitory weights adjust in order to reach an approximation of a detailed balance (Figure 4E, F). Because of the broader input tuning in the inhibition, the stimulus tuning of the inhibitory currents remains broader than that of the excitatory current (Figure 4F), similar to physiological findings in primary visual cortex (Liu et al., 2011). Although the final tuning of the excitatory and inhibitory input currents is relative wide, the resulting firing rate tuning of the cell (i.e., the rectified difference of the excitatory and inhibitory currents) is relatively narrow, due to an ‘iceberg’ effect in which excitation supersedes inhibition only in a small stimulus range (Figure 4F, dashed line).

**Figure 4:**
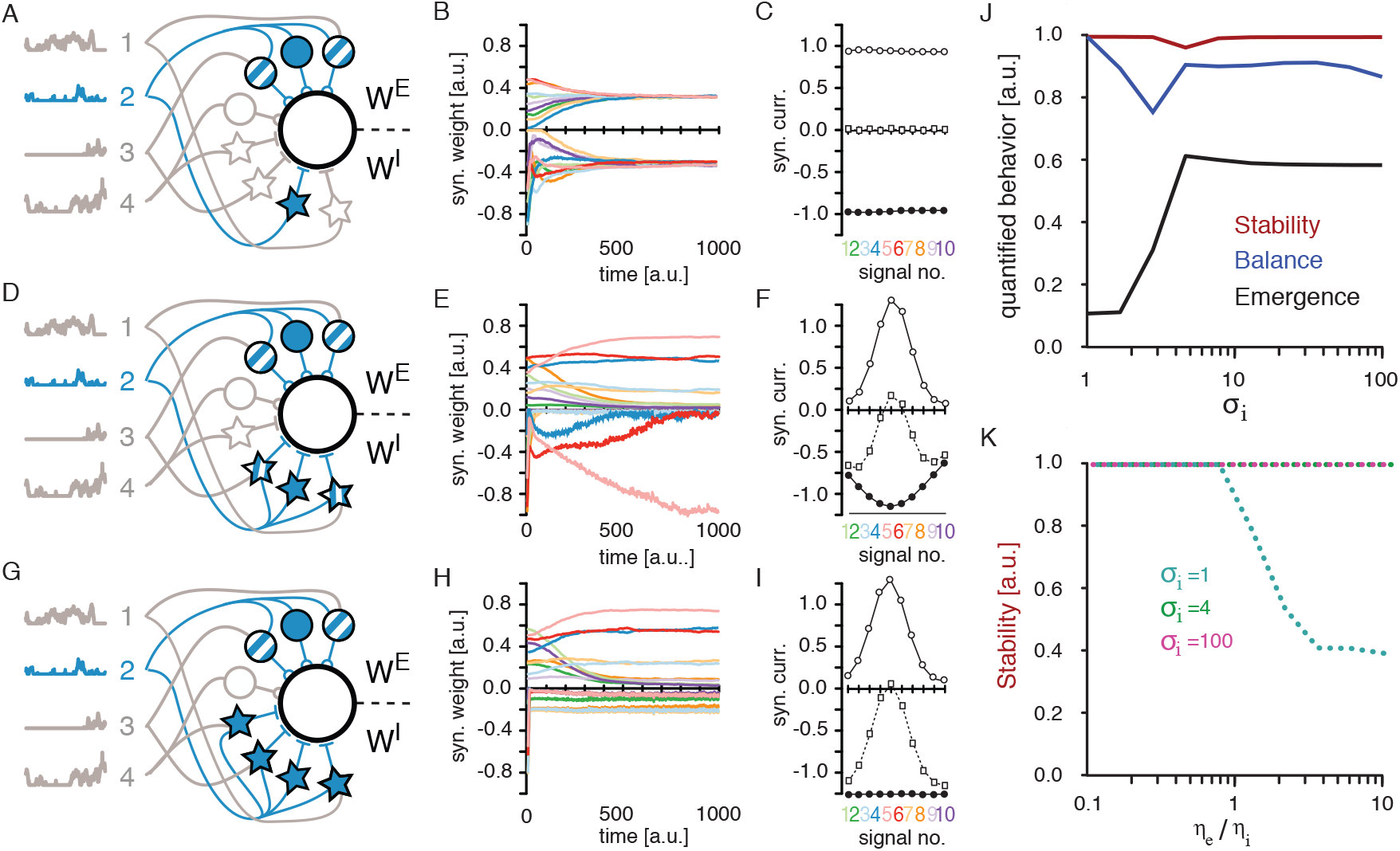
Broader inhibitory than excitatory tuning supports receptive field formation. A) Network diagram. A single postsynaptic neuron receives synaptic inputs from excitatory (circles) and inhibitory (stars) populations. Each input population fires according to a weighted superposition of the input signals, with weights that follow a Gaussian distribution. Effectively, this introduces a Gaussian tuning of the input populations as a function of input signal. The tuning width of the excitatory inputs was kept constant (*σ_E_* = 1), while the tuning width of the inhibitory inputs was systematically varied (B,C: *σ_I_* = 1; E,F: *σ_I_* = 4; H, I: *σ_I_* = 100). B, E & H) Temporal evolution of excitatory (positive) and inhibitory (negative) synaptic weights. C, F & I) Synaptic currents after learning (excitation: open circles, inhibition: filled circles, net current: open squares). J) Dependence of stability, balance and emergence indices on the relative tuning width *σ_E_*/*σ_I_* of excitation and inhibition. K) Dependence of stability on the relative learning rates of excitation and inhibition for different inhibitory tuning widths *σ_I_*.

Increases in inhibitory tuning width have only a minor impact on the stability of the final synaptic configuration, but introduce a relatively sharp transition to the emergence of a receptive field at the cost of a reduced precision of the E/I balance (Figure 4J). At the transition point, both the E/I balance and the stability is slightly reduced, because the homogenous weight configuration loses stability through an oscillatory bifurcation, i.e., the synaptic weights oscillate in a small range of inhibitory tuning widths around the transition point (not shown). When the inhibitory input tuning is wider than the excitatory tuning, the output neuron can under certain conditions develop a periodic tuning with respect to the input channel, reminiscent of periodic receptive fields that have been found in the hippocampal formation (Hafting et al., 2005). Finally, broadened inhibition removes the oscillatory instability that was observed for high excitatory learning rate for stimulus-specific inhibition (Figure 4K).

### Co-tuned receptive fields can emerge from a competitive normalization and input inhomogeneities

It is well-known that different normalization schemes for the excitatory weights support the emergence of stimulus selectivity to a different degree (Miller & MacKay, 1994). In particular, a subtractive normalization gives rise to stronger competition between synapses than the multiplicative normalization we have used to far (Dayan & Abbott, 2001). However, with subtractive normalization we observed qualitative differences only for specific inhibition and rapid inhibitory plasticity. In this case, the excitatory weights don’t converge to equal strength (cf. Figure 2), but perform a random walk that generates an unstructured, temporally fluctuating receptive field (Figure 5A, B). These dynamics arise because, on average, the effective learning rule Eq. 3 introduces weight changes that are immediately reverted by the normalization (Figure 5C), such that all possible weight configuration are marginally stable. Changes in the relative learning rates of excitatory plasticity and the relative tuning widths of the excitatory and inhibitory inputs had qualitatively similar effects as for the multiplicative normalization (Figure 5D, E).

**Figure 5:**
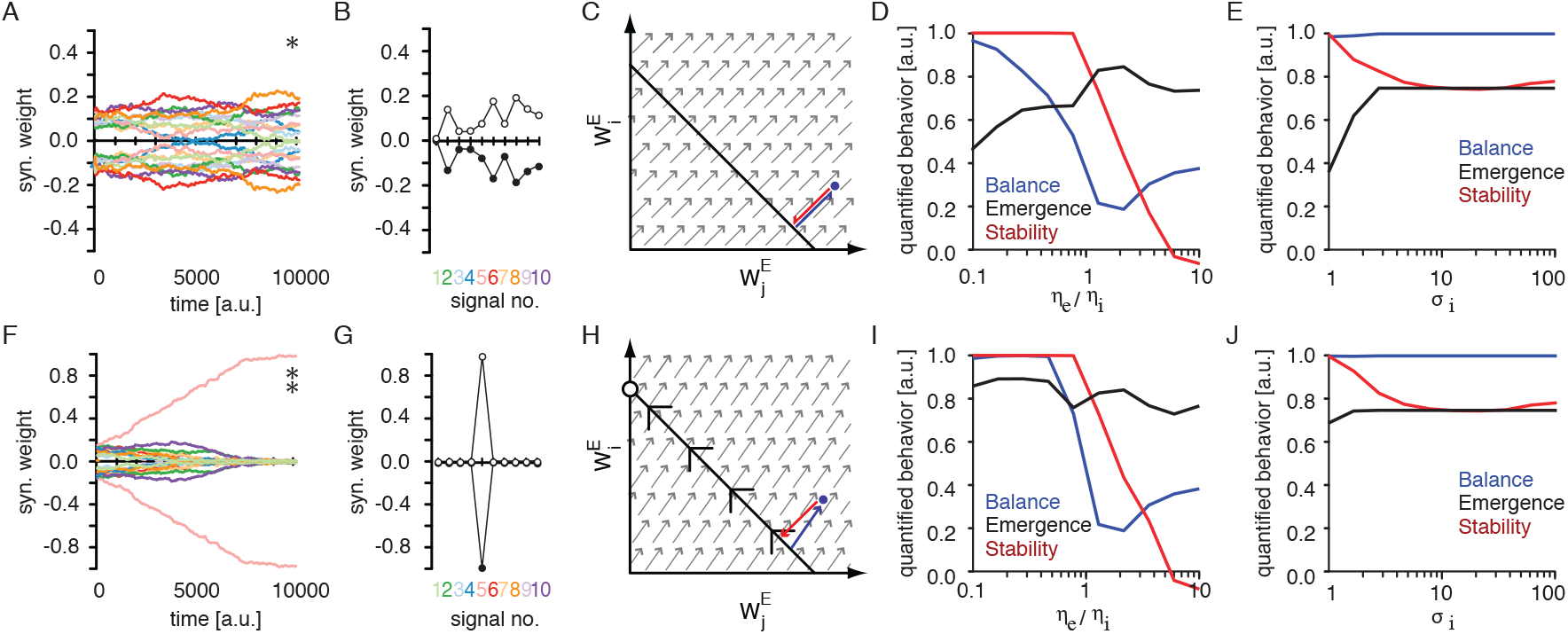
Co-tuned receptive fields for subtractive normalization and biased inputs. A) Temporal evolution of excitatory (positive) and inhibitory (negative) synaptic weights. B) Synaptic weights after learning. C) Illustration (for two excitatory weights only) of the mechanism that governs the dynamics in the synaptic weights. As in Figure 3, the excitatory rule aims to increase all excitatory weights by the same amount (C, grey arrows and blue arrow). On average, these changes are now exactly counteracted by the subtractive normalization that reduces all weights by the same amount (red arrow). As a result, the whole constraint line is marginally stable (black line), and the weight dynamics are dominated by fluctuations. D, E) Dependence of stability, balance and emergence indices on the relative learning rate *η_E_*/*η_I_* (D) and the relative tuning width *σ_E_*/*σ_I_* (E) of excitation and inhibition. F-J) same as A-E, but the activity of input signal 5 is increased by 10%. H) The excitatory learning rule now causes more potentiation for the weights of one population (grey arrows, blue arrow for an example of a Hebbian weight update), which in combination with the subtractive normalization (red arrow) leads to a full specialization of the neuron for input population 5 (black circle). This strong specialization is not present for a multiplicative normalization (Fig. 5 - Supplement 1)

A mathematical analysis (Eq. 3 and SOM) suggests that this random walk behavior requires that all excitatory inputs have exactly the same mean firing rates. If one excitatory input has a higher mean firing rate than the others (+10% in our simulations), its weights will increase more rapidly, leading to a “tilt” in the vector field (Figure 5H). The synaptic weight of the input with the highest firing rate will thus outgrow all others, leading to the formation of a stimulus-selective excitatory receptive field that is balanced by precisely co-tuned inhibition (Figure 5F, G). The increased firing rate of one input did not change the dependence of the dynamics on the relative learning rates of excitatory and inhibitory plasticity (Figure 5I) or the relative tuning widths of excitation and inhibition (Figure 5J).

In summary, the interaction of excitatory and inhibitory plasticity, combined with a competitive weight normalization, can amplify small input inhomogeneities and lead to the development of receptive fields with a precise co-tuning of excitation and inhibition, similar to the receptive fields that are found in auditory cortex (A1). We therefore studied whether the model can also reproduce other dynamical phenomena that are observed during receptive field plasticity in A1 (Froemke et al., 2007).

### Stimulus-selective inhibition with subtractive normalization explains auditory receptive field shape and plasticity

Neurons in primary auditory cortex often have bell-shaped tuning curves with respect to the frequency of pure tones, both in terms of their firing rate and their excitatory and inhibitory input currents (Wehr & Zador, 2003). Their excitatory and inhibitory tuning functions are often co-tuned, an effect that gets more pronounced during development and seems to be driven by sensory experience (Figure 6A) (Dorrn et al., 2010). Moreover, in adult animals, Froemke et al. (2007) have shown that both excitatory and inhibitory tuning functions remain stable in the presence of pure tone stimulation (Figure 6B) unless the tones are paired with neuromodulatory input, e.g., from nucleus basalis (NB), the main source of cortical acetylcholine (Figure 6C). In response to paired NB and pure tone stimulation, the excitatory tuning curve of A1 neurons shifts its maximum (i.e., its preferred frequency) to that of the presented tone (Figure 6C, middle). This stimulation paradigm initially leaves the inhibitory tuning unchanged. However, in the presence of auditory stimulation the inhibitory tuning curve gradually shifts to the new preferred frequency of excitation, until a new state of co-tuning is reached after a few hours (Figure 6C, right). Interestingly, over even longer periods, both the excitatory and the inhibitory tuning curves revert back the original preferred frequency (Figure 6D).

**Figure 6:**
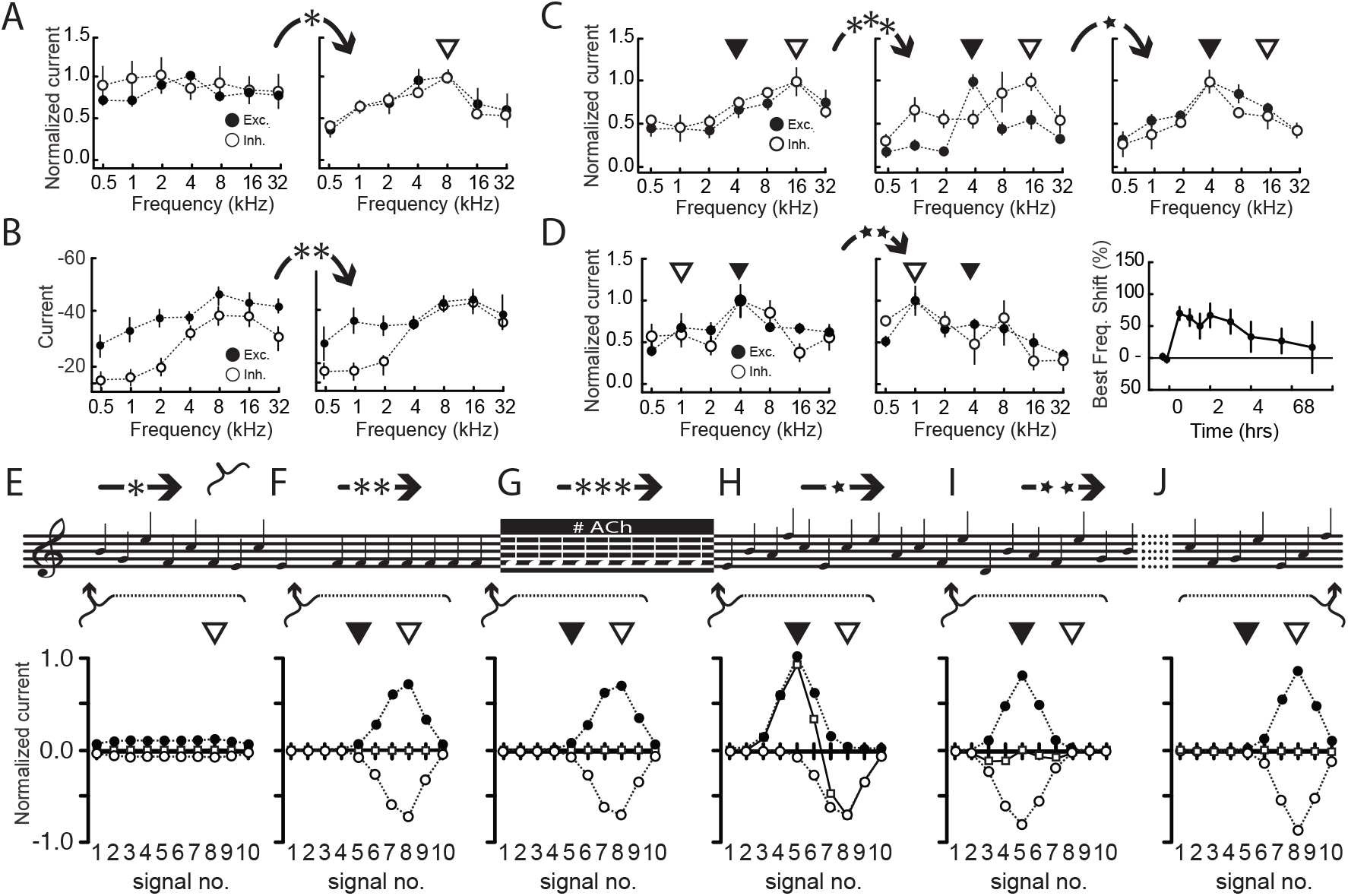
Interacting excitatory and inhibitory synaptic plasticity reproduce receptive field plasticity. A-D) Electrophysiological data from rat primary auditory cortex (A1). A) Co-tuning of excitatory and inhibitory receptive fields in A1 increases during development. Imbalanced synaptic frequency tuning at P14 (left), balanced frequency tuning in adult rats (right). Filled circles, excitation; open circles, inhibition. B) In adults, receptive fields are robust to pure tone stimulation alone. Normalized frequency tuning of excitation and inhibition before stimulation (left) and after stimulation (right). C) Excitatory receptive field plasticity induced by paired pure tone and nucleus basalis stimulation. Tuning curves before (left) and after paired stimulation (middle). Inhibitory receptive fields shift to rebalance excitation within hours (right). D) Duration of synaptic frequency tuning modifications induced by a single episode of nucleus basalis pairing. Left, normalized frequency tuning curves for an A1 neuron recorded 125 minutes after pairing. Middle, a different cell from same A1 region recorded 295 minutes after pairing. Right, time course for normalized shift in excitatory tuning curve peak. 0% represents the original best frequency for a given A1 location; 100% is a full shift to the paired frequency. Data from 52 recordings in 24 animals. Time is relative to the end of pairing. Error bars show s.e.m.. Data with permission from Dorrn et al. (2010) (A), Froemke et al. (2007) (B, C) and Martins & Froemke (2015) (D). E-J) Computational model. E) Synaptic weights are initialized to a weak excitatory and inhibitory stimulus tuning. Stimulus channel 8 (preferred channel - open triangle) is 10% stronger than the other channels. F) Sensory stimulation causes an emergence of co-tuned and bell-shaped excitatory and inhibitory tuning curves at the preferred channel no 8. G) In the balanced configuration, pure tone stimulation (at stimulus channel no 5) causes only minor changes of excitatory and inhibitory tuning curves. H) Pairing pure tone stimulation with disinhibition shifts the maximum of the excitatory tuning curve to the frequency of the pure tone (training channel no 5 - full triangle). Inhibitory tuning remains largely unchanged. I) Inhibitory synaptic plasticity triggered by sensory experience shifts the inhibitory tuning to rebalance excitation (peak at training channel no 5). J) Extended sensory experience shifts both excitatory and inhibitory receptive fields back to their original preferred channel (#8).

To investigate whether an interaction of excitatory and inhibitory synaptic plasticity can reproduce these rich dynamics of receptive field plasticity, we interpreted the different input channels as auditory frequencies. Again, one of the excitatory inputs has a higher firing rate and the excitatory weights are subject to a subtractive normalization. Under these conditions, the interplay of excitatory and inhibitory plasticity in the presence of sensory stimulation leads to the development of bell-shaped tuning curves for both excitatory and inhibitory currents, peaking at the same input channel (Figure 6E→F). After this co-tuning has been established, stimulation of an individual input channel causes only small changes in both excitatory and inhibitory tuning curves (Figure 6F→G), as observed in the experiment. This stability arises from the detailed balance established by the inhibitory plasticity that leads to low firing rates close to the target rate *ρ*_0_, and from small learning rates. It has been hypothesized that cholinergic inputs as evoked by NB stimulation cause a transient disinhibition of cortical pyramical cells (Froemke et al., 2007; Letzkus et al., 2011). Hence, we mimicked NB stimulation by a transient suppression of the firing rate of the inhibitory inputs. Pairing such an “NB stimulation” with the activation of a non-preferred input channel shifts the peak of the excitatory tuning curve to the stimulated input channel, while leaving the inhibitory tuning curve un-altered (Figure 6G→H). The shift in the excitatory tuning curve is caused by high postsynaptic firing rates during disinhibition, while inhibitory plasticity is reduced by the small firing rates of the inhibitory input neurons. Subsequent random stimulation of all input channels causes the same gradual re-balancing dynamics that are observed in A1 (Figure 6H→I)(Vogels et al., 2011). On an even longer time scale, both the excitatory and inhibitory tuning curves slowly revert back to the original preferred frequency, due to the higher firing rate of the corresponding input channel (Figure 6I→J). In summary, the interaction of excitatory and inhibitory Hebbian plasticity seems to be sufficient to reproduce the rich dynamics of receptive field plasticity in A1.

## Discussion

Our analysis suggests that concurrent excitatory and inhibitory Hebbian plasticity can generate a rich repertoire of receptive field dynamics. In particular, we identified two essential factors that control their interaction: The stimulus-specificity of the inhibitory inputs and the relative degree of plasticity of excitatory and inhibitory synapses. Unspecific, but plastic feedforward inhibition generates a sliding threshold for neuronal activity that leads to the formation of a receptive field with high stimulus-selectivity, by a mechanism that is similar to that of BCM rules (Bienenstock et al., 1982). This observation could be relevant in the context of the search for a biophysical basis of the sliding threshold of BCM theory (Cooper & Bear, 2012). In place of a direct dependence of the excitatory plasticity rule on previous activity, our analysis suggests that a sliding threshold can be implemented indirectly, by adaptive inhibition that changes how a postsynaptic neuron responds to a given excitatory input (Miller, 1996; Triesch, 2007).

Our analysis also suggests that for stimulus-specific inhibitory inputs, the homeostatic action of the inhibitory plasticity rule applied here (Vogels et al., 2011) equilibrates the firing rates to different stimuli and therefore does not favor a spontaneous formation of stimulus selectivity. This democratic tendency can be broken in different ways. Increases in the tuning width of the inhibitory inputs favor the formation of a receptive field, at the cost of a less precise E/I co-tuning. Alternatively, receptive field formation can be promoted by competitive weight limiting mechanisms (such as subtractive normalization), which can amplify slight asymmetries in the input statistics.

The model further supports the hypothesis that disruptions of the E/I balance (specifically, transient increases of excitation or decreases of inhibition) could serve as a gate for the induction of plasticity. This idea is in line with observations that the E/I balance is less precise in young animals (Dorrn et al., 2010), with the hypothesis that the maturation of inhibition controls the duration of developmental critical periods (Hensch, 2005; Kuhlman et al., 2013) and with the apparent need for disinhibition for receptive field plasticity in mature animals (Froemke et al., 2007; Kuhlman et al., 2013). In our simulations, the detailed E/I balance that is established by inhibitory plasticity provides a default state in which the gate for the induction of synaptic plasticity is closed. Perturbations of this balance, e.g., by selective disinhibition, open the gate. Of course, our model captures only one aspect of the wide range of neuromodulatory effects. Neuromodulators are likely to influence synaptic plasticity through other pathways, e.g., by directly affecting the biophysical machinery of synaptic plasticity (Pawlak et al., 2010) or the electrical properties of neuronal arborizations (Tsubokawa & Ross, 1997; Wilmes et al., 2016).

We concentrated our analysis on a relatively simple and largely linear model that is amenable to mathematical analysis. However, we expect that many of the dynamical phenomena will generalise to other neuron models and learning rules. We have observed similar dynamics when the excitatory learning rule was replaced by a rate-based triplet rule that includes long-term depression (Pfister & Gerstner, 2006; Clopath et al., 2010), as long as the threshold rate between potentiation and depression in the triplet rule was lower than the target rate of the inhibitory plasticity. If this threshold was higher than the target rate, however, all excitatory weights converged to zero. For rapid inhibitory plasticity, the same dynamics are observed for the interaction of the inhibitory plasticity rule with classical BCM rules (Bienenstock et al., 1982), because the rate homeostasis of the inhibitory plasticity keeps the sliding threshold of the BCM rule largely constant, thereby effectively reducing the BCM rule to a triplet rule without sliding threshold. Because nonlinearities in the learning rule are closely related to nonlinearities in the neuronal transfer function, we also expect a qualitatively similar behavior for nonlinear neuron models. From a more abstract perspective, the presently analyzed model can be interpreted as a linearization around a homogenous state in which all excitatory and inhibitory weights are equal. Our results on the stability of this state will apply locally and provide an indication of whether a neuron will develop a receptive field or remain unselective.

Our analysis is limited to feedforward networks, although the co-tuning of excitatory and inhibitory inputs in early sensory areas could in principle arise from either stimulus-selective feedforward inhibition or feedback inhibition. Feedback inhibition appears as a natural candidate to explain the observed E/I co-tuning in cortical areas with a topographical organization (Harris & Mrsic-Flogel, 2013). However, the dissociation of the excitatory and inhibitory tuning curves induced by Froemke et al. (2007) indicates that a component of stimulus-selective feedforward inhibition is present in auditory cortex. We suspect that our results could generalize to network architectures with recurrent inhibition, provided that a sufficiently rich pool of sensory tuning curves is present in the inhibitory population. However, an analysis of receptive field dynamics in recurrent networks is considerably more difficult, because both forms of synaptic plasticity would change not only the postsynaptic tuning function, but also that of the inhibitory inputs.

To limit the complexity of the system, we ignored temporal aspects of neuronal and synaptic integration as well as the spike timing dependence of synaptic plasticity. In particular, inhibitory synapses tend to have slower dynamics than excitatory synapses (Wehr & Zador, 2003). Rapid stimulus transients therefore cannot be balanced and cause reliably timed onset spikes (Vogels et al., 2011). Synaptic dynamics thus impose limits on the precision of the E/I balance that can be reached by inhibitory plasticity. In combination with excitatory spike timing-dependent plasticity, this is likely to introduce a selectivity of the neuron to temporal input features (Kleberg et al., 2014; Sterling & Sprekeler, 2014).

Unfortunately, the experimental characterization of inhibitory synaptic plasticity is less advanced than that of excitatory plasticity, and earlier work has drawn a somewhat diverse picture (Woodin & Maffei, 2010; Vogels et al., 2013). In part, this is likely due to the diversity of inhibitory cell types (Markram et al., 2004; Klausberger & Somogyi, 2008; DeFelipe et al., 2013) and their largely unresolved functional roles. However, a recent study of inhibitory plasticity in auditory cortex supports our core assumption that Hebbian inhibitory plasticity aids in establishing an E/I balance (D’amour & Froemke, 2015; Kirkwood, 2015), and parvalbumin-positive interneurons are emerging as potential mediators within the cortical microcircuit (Xue et al., 2014). We tested a few variants of the inhibitory learning rule. As long as the rule remained Hebbian, i.e., coincident pre- and postsynaptic activity predominantly caused potentiation, and inhibitory weights decayed in the absence of postsynaptic activity, the rule established an approximate balance of excitation and inhibition (Luz & Shamir, 2012). However, the democratic aspect of the presently studied rule may be less pronounced for other rules, potentially facilitating the emergence of a receptive field even for specific inhibition.

## Conclusion

Our study provides a theoretical underpinning for the joint emergence of excitatory and inhibitory receptive fields in sensory cortices and adds a developmental aspect to the discussion of how the stimulus specificity of interneurons is related to the tuning properties of excitatory cells (Harris & Mrsic-Flogel, 2013). What’s more, it provides a mechanistic description of the experimentally observed gating of experience-dependent plasticity in adult animals by disinhibitory and neuromodulatory mechanisms.

## Experimental Procedures

### Neuron model and network structure

We study a feedforward network consisting of a single neuron receiving both excitatory and inhibitory inputs. To keep the system simple and allow an analytical treatment of the learning dynamics, we study a threshold-linear neuron and concentrate on a rate-based description of neural activity. When we refer to input activity or synaptic weights, we thus mean firing rates of neural populations and total synaptic connection strengths between input populations and the output neuron. Given time-dependent activities *E_i_*(*t*) and *I_j_*(*t*) of *N_E_* excitatory and *N_I_* inhibitory inputs, respectively, the output rate of the model neuron is given by

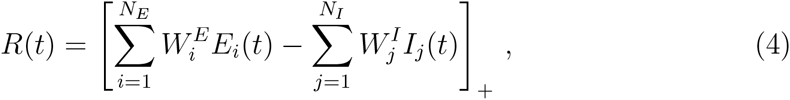

where 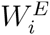 and 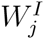 denote the synaptic weights of the excitatory and inhibitory synapses, respectively, and [·]_+_ denotes a rectification that sets negative values to zero, to avoid negative firing rates. To comply with the notion of excitation and inhibition, all synaptic weights are constrained to be positive. In all simulations, we model *N_E_* = 10 excitatory input populations. For the simulations with unspecific inhibition, the neuron receives a single inhibitory input *N_I_* = 1, in all other simulations there are as many excitatory as inhibitory input channels: *N_I_* = *N_E_* = 10. Note that this is a statement about how many functionally different populations of inhibitory neurons project to the output cell, not about the number of presynaptic cells, which will in general be different for excitation and inhibition.

### Input signals

The excitatory and inhibitory input signals *E_i_* and *I_j_* are generated assuming that the inputs each have a tuning to sensory stimuli. These stimuli are modeled as *N* = 10 different sensory stimulus channels with time-dependent activities *s_j_*(*t*) (which could, e.g., be sound amplitude at different frequencies). The activity of input neuron *i* is calculated by a sum of the stimulus channels, weighted with tuning strengths 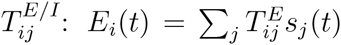 and 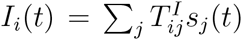. The input tuning is Gaussian: 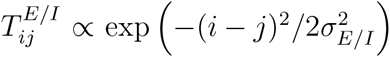 and normalized such that Σ_*j*_ *T_ij_* = 1 for all *i*. The parameters *σ_E_*/_*I*_ denote the tuning widths for excitation and inhibition, respectively. In the limit of very small tuning width, the input signals are exact copies of the activity in the sensory stimulus channels; for very large tuning widths, they are an average thereof.

The activities *s_i_*(*t*) of the stimulus channels are generated from independent Ornstein-Uhlenbeck processes with a time constant of 50 ms by subtracting a constant *c*, setting all negative values to zero and then rescaling the signal to have a mean firing rate of 1 (arbitrary units). The constant c controls the lifetime sparseness (Franco et al., 2007) of the signals. For our choice of *c*, we obtained a lifetime sparseness of 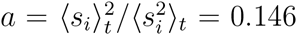, where 〈·〉_*t*_ denotes a temporal average. The results are robust to the precise value of the sparseness of the input signals, which mainly controls how well the output neuron can differentiate between the input signals. It thereby indirectly controls the convergence of the learning dynamics. The sparser the input signals, the higher the learning rate can be chosen before the dynamics become obstructed by the noisy dynamics of the online learning rule.

In the case of unspecific inhibition, we simulate a single inhibitory input channel, the activity of which is constant in time. The dynamics do not change when, instead, the inhibitory input is an average of the activities of the excitatory inputs (i.e., 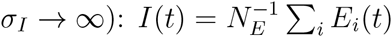 (see SOM for mathematical analysis).

In simulations where the subtractive normalization amplifies differences in input firing rates (Figs. 5 and 6), one of the stimulus channels *s_j_* was multiplicatively scaled up by 10%.

### Excitatory synaptic plasticity

We study a simple Hebbian learning rule for the excitatory synapses

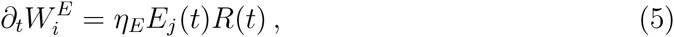

where *η_E_* denotes the excitatory learning rate, which is 10 times smaller than the inhibitory learning rate, unless specified otherwise. Excitatory plasticity is inherently unstable, so this rule has to be complemented by a weight-limiting mechanism. Because previous work has shown that the specific form of the weight limiting mechanism is important for the learning dynamics (Miller & MacKay, 1994), we study both multiplicative and subtractive weight normalization. A multiplicative normalization is vaguely inspired by homeostatic synaptic scaling (Turrigiano et al., 1998). Note that an *activity*-dependent homeostatic control of the excitatory synaptic weights (rather than a weight-dependent mechanism as used here) is problematic in a situation where inhibitory synapses are also plastic, because neuronal activity and excitatory weights are only weakly coupled. For example, both excitatory and inhibitory weights could diverge although a given mean firing rate is maintained.

In our simulations, we start with random weights drawn from a uniform distribution. For multiplicative normalization, after every weight update, the weights are divided by their *L*_2_ norm. For the subtractive weight normalization, we subtract the average weight 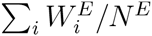 from all weights and add a constant (here 1). Negative weights, which can arise from this procedure, were clipped to zero. By this procedure, the sum of the weights remains at approximately *N^E^*.

### Inhibitory synaptic plasticity

The inhibitory synapses of the network are plastic according to the balancing learning rule we previously suggested (Vogels et al., 2011)

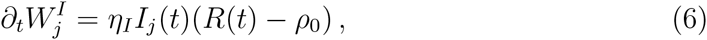

where *η_I_* is the inhibitory learning rate (Figs. 1–4: *η_I_* = 10^−3^, Figs. 5, 6: *η_I_* = 10^−2^). The learning rule Eq. 6 introduces a homeostatic control of the firing rate: inhibitory synaptic weights are adjusted such that the output rate of the neuron approaches a target rate *ρ*_0_. If the excitatory currents received by the neuron are large (i.e., if the activity of the neuron due to the excitatory input alone is large compared to the target rate *ρ*_0_), excitatory and inhibitory input currents to the neuron become approximately balanced, with a precision that is determined by the correlation between excitatory and inhibitory input currents (Vogels et al., 2011). In all simulations, the target rate was *ρ*_0_ = 0.01 (again in arbitrary units).

### Disinhibition by neuromodulation

In the last figure, we study the effect of neuromodulation on learning, with the goal of reproducing the experimental data of Froemke et al. (Froemke et al., 2007). To this end, we first develop balanced receptive fields with a preference for signal number 8, by increasing the input for signal number 8 by 10% (i.e., an average firing rate of 1.1 instead of 1). The excitatory learning rule is paired with a subtractive normalization, and *σ_E_* = *σ_I_* = 1. During training, only signal number 5 is active at constant rate of 5, mimicking the presentation of a pure tone. Finally, when training is paired with acetylcholine, we reduce feedforward inhibition as shown experimentally (Xiang et al., 1998; Letzkus et al., 2011) by setting the tuning strengths *T^I^* to 0.

### Network quantification

We quantify the network by three different parameters, the stability, the balance and the emergence of receptive fields. The stability *S* is the average inner product of the excitatory weights (L2-normalized) at two different time points, averaged over 1000 random samples. These time points are chosen randomly in the second half of the simulation to insure learning convergence. The balance *B* is average correlation between excitatory and the inhibitory weights (rescaled so that the maximum weight is one) over the second half of the simulations. Finally, the emergence *E* is computed as one minus the ratio of the mean excitatory weights and their current maximum, the ratio is then averaged over the second half of the simulation.

## Acknowledgments

HS is supported by the German ministry for Science and Education (grant no. 01GQ1201) and performed parts of this research at the University of Cambridge, UK, and the Humboldt-Universität zu Berlin. RCF is supported by NIDCD grants DC12557 and DC009635, a Sloan Scholarship and a Klingenstein Fellowship. TPV was supported by a Sir Henry Dale Wellcome Trust and Royal Society Fellowship (WT100000). We would like to thank Simon Weber for comments on the manuscript.

## Figures & Figure legends

**Figure 1 - Supplement 1:**
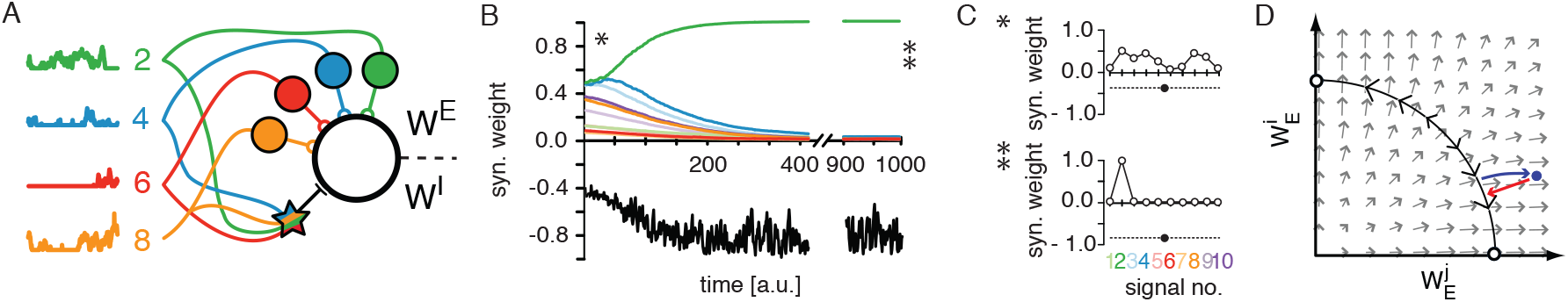
Receptive field formation with pooled feedforward inhibition. A) Network diagram. A single postsynaptic neuron receives synaptic input from 10 excitatory populations (colored circles, not all 10 shown) with different time-varying firing rates (signals, colored traces) and from a single inhibitory population, which pools over the 10 excitatory inputs (grey star). B) Temporal evolution of excitatory (positive) and inhibitory (negative) synaptic weights. C) Synaptic weights before (top) and after learning (bottom). Stars indicate corresponding times in B. D) Illustration of the mechanism that leads to the emergence of selectivity in the synaptic weights. Parameters: excitatory learning rate *η_E_* = 10^−3^, inhibitory learning rate *η_I_* = 10^−4^, target rate of inhibitory plasticity *ρ*_0_ = 10^−2^.

**Figure 5 - Supplement 1:**
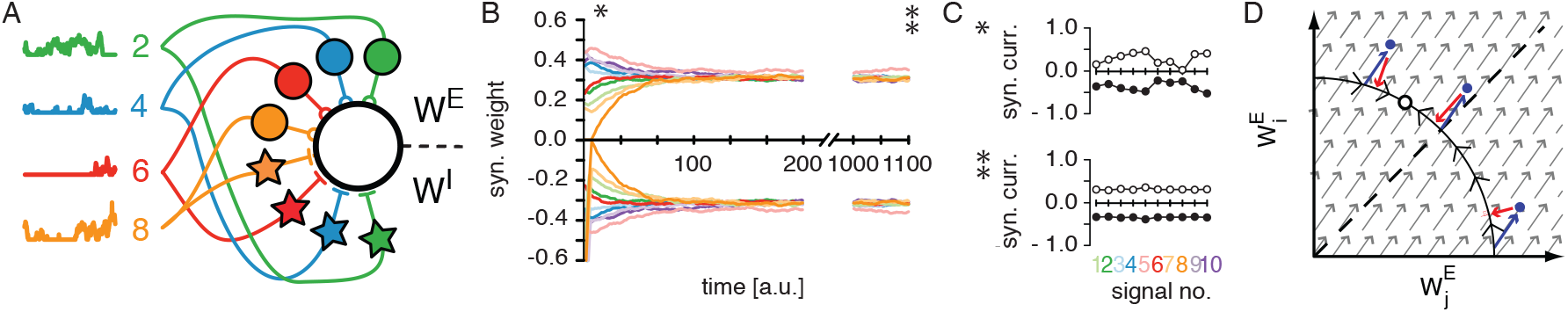
Asymmetries in presynaptic firing rates have gradual effects for multiplicative normalization. A) Network diagram. A single postsynaptic neuron receives synaptic input from 10 excitatory populations (colored circles, not all 10 shown) with different time-varying firing rates (signals, colored traces) and input from 10 corresponding inhibitory populations with the same rates. The average activity for signal 5 was increased by 10% compared to the other signals. B) Temporal evolution of excitatory (positive) and inhibitory (negative) synaptic weights. C) Synaptic weights before (top) and after learning (bottom). Stars indicate corresponding times in B. After learning, the synaptic weight for input signal 5 is only mildly higher than those of the other signals. D) Illustration of the mechanism that causes the gradual dependence on the mean presynaptic firing rate. Parameters as in Figure 5, apart from normalization.

## Supplementary Online Material for

### 1 Derivation of effective learning rules

In the case where the inhibitory learning rate is much higher than the excitatory learning rate, *η_I_* ≫ *η_E_*, the excitatory learning rule can be replaced by an effective learning rule that incorporates the effects of plastic inhibition. To derive such effective learning rules, we consider two cases: the case of specific inhibition and the case of unspecific inhibition.

#### Unspecific inhibition

For unspecific inhibition, we assume that all inhibitory inputs have the same, albeit potentially time-varying firing rate, so that we can reduce the problem to one single inhibitory input with firing rate *I*. We study two cases: In the first, the inhibitory input firing rate is constant (*I*(*t*) = *I* = *const*.), in the second, it is given by the population activity of the excitatory inputs (*I*(*t*) = Σ_i_ *E_i_*(*t*)). The former case corresponds to uncorrelated tonic inhibition, the latter to pooled feedforward inhibition.

For uncorrelated tonic inhibition, we can insert the expression for the postsynaptic firing rate *R*(*t*) into the Hebbian learning rule to obtain the effective excitatory learning rule stated in the main text:

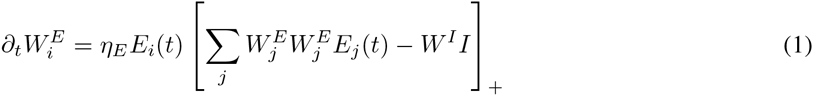

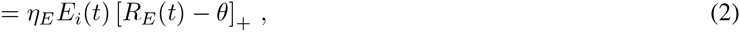

where 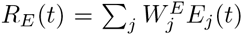 is the total excitatory input to the cell and *θ* = *W^I^ I* the inhibitory input. Structurally, this rule is similar to a BCM rule with θ acting as a threshold. To show that the threshold is sliding, we only need to consider the inhibitory learning rule, multiplied with the inhibitory input rate *I*:

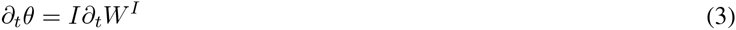

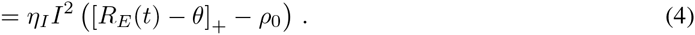

For sufficiently small learning rates (such that we can consider the time-averaged version of the learning dynamics), the stable fixed point of this equation is given by the implicit condition

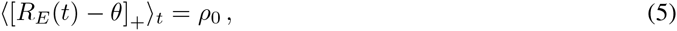

where 〈·〉_*t*_ denotes temporal averaging. The speed at which the threshold converges to this fixed point is mainly determined by *η_I_I*^2^, i.e., by the inhibitory learning rate and the firing rate of the inhibitory inputs.

The case where the inhibitory firing rate *I*(*t*) is given by the mean activity *Ē*(*t*) = *N*_−1_ Σ_*i*_*E*(*t*) of the excitatory rates 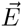 creates a slightly different situation, because the inhibitory input depends on the excitatory input and hence varies in time. The stationary state for the inhibitory weight *W^I^* (again assuming sufficiently small learning rates) is determined by the equation

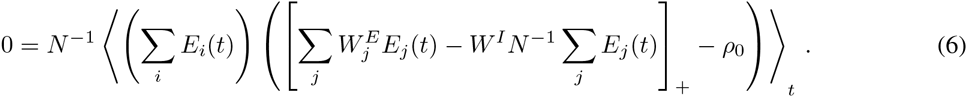

To find an analytical solution that can be understood intuitively, we neglect the output rectification in the learning dynamics of the inhibition (admittedly a rather questionable approximation), and rewrite equation 6 in vector notation using the covariance matrix 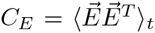 of the excitatory inputs and the homogeneous weight vector 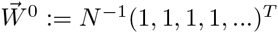:

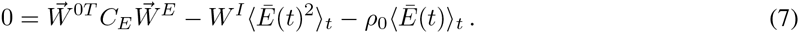

If we assume that the statistics of the excitatory inputs are symmetric in the sense that the homogeneous vector 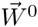 is an eigenvector of the excitatory covariance matrix *C_E_*, we can calculate an explicit expression for the stationary inhibitory weight:

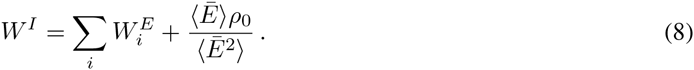

If we insert the resulting output firing rate

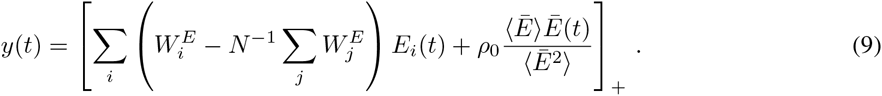

into the excitatory learning rule, we also get a Hebbian rule with a “sliding threshold”, but the threshold is not given by the temporal average of the excitatory drive, but by the momentary excitatory drive that would be caused by a homogeneous weight vector of the same total synaptic weight. From this perspective, this rule generates a spatial competition between synapses, while the case of tonic inhibition generates a temporal competition between stimuli. Both lead to the formation of a receptive field, as shown in Fig. 1 of the main text and the associated Supplement 1.

The validity of the effective learning rule resulting from Equation 9 is questionable, because the derivation first neglects the output rectification of the neuron and later reintroduces it, but it nevertheless provides an intuition for the mechanism behind the symmetry breaking observed in the simulations.

#### Specific inhibition

By specific inhibition we mean that the inhibitory inputs contain a sufficient stimulus selectivity that a balance of excitatory and inhibitory inputs can be reached on a moment-by-moment basis, for arbitrary excitatory weights *W^E^*. To ensure this, it is sufficient and necessary in the present linear picture that all excitatory inputs can be written as a linear combination of the inhibitory inputs, i.e., that there is a matrix *M* such that

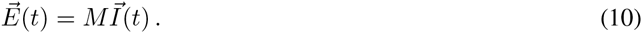

The stationarity condition for the inhibitory weights is

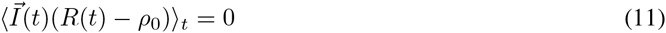

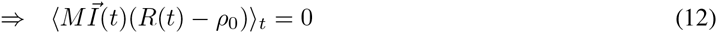

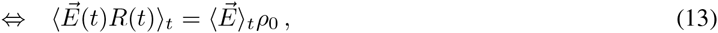

which can be directly inserted into the averaged weight dynamics of the excitatory weights:

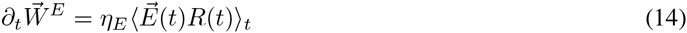

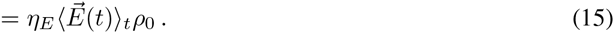

By taking the online version of this learning rule and reverting back to index notation, we get the the effective learning rule that was intuitively motivated in the main text:

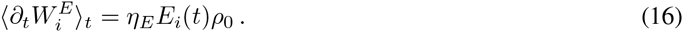

### 2 Mathematical analysis of the learning dynamics for specific inhibition

To study the properties of the fixed points of the full system of excitatory and inhibitory plasticity, we need to take the effects of the normalization into account. As shown by Miller and MacKay (1994), both a multiplicative and an subtractive normalization can be included in a dynamical system by an additional normalization-specific term in the excitatory learning rule:

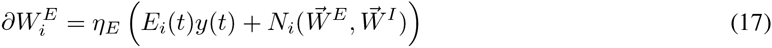

with 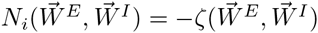 independent of *i* for the subtractive normalization and 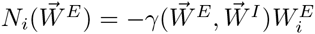 for the multiplicative normalization. Here, *γ* is a scalar function of the excitatory weight vector that is independent of *i*. The specific shape of the functions *ζ* and *γ* controls the shape of the constraint manifold.

#### Fixed points

To find the fixed points of the coupled learning rules for excitation and inhibition, we can first find the fixed points of the inhibitory learning rule and insert it into the excitatory rule. In the case where the inhibition is specific, the calculation of the fixed points of the excitatory weights thus amounts to finding the fixed points of the effective learning rule Eq. 15, enriched by the additional constraint terms. For an subtractive normalization, this leads to the fixed point equation:

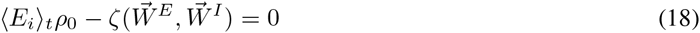

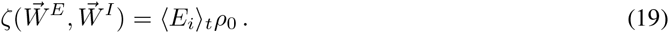

If the input statistics are the same, i.e. all 〈*E_i_*〉_*t*_ have the same value, this reduces to a single equation, suggesting that any point on the constraint manifold for the excitatory weights is a fixed point. This is in line with the diffusive dynamics observed in the simulations. If the statistics are not the same, this equation has no solution, suggesting that the fixed point(s) will lie at the border of the constraint manifold. Small differences in the mean input firing rates thus have a drastic effect, as observed in the simulations in Fig. 5 of the main text.

For a multiplicative normalization, the fixed point equation has the following form

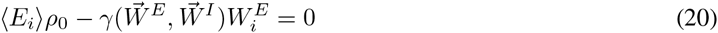

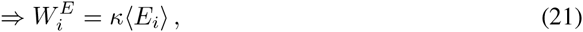

where *κ* is a constant that has to be chosen such that the normalization requirement is fulfilled. If the mean firing rate 〈*E_i_*〉 is the same for all input neurons, the only fixed point is the homogenous solution in which all excitatory synapses have the same strength, in agreement with the simulation results. Moreover, small differences in the mean firing rate of the excitatory inputs lead to gradual changes of the fixed point, in contrast to the drastic impact they have for an subtractive normalization (main text Fig. 5 - Supplement 1).

#### Jacobian at the fixed points

To evaluate the stability of the fixed points, we have to calculate the Jacobian of the learning dynamics. For specific inhibition (i.e., 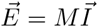) and sufficiently small *ρ*_0_, the Jacobian is given by

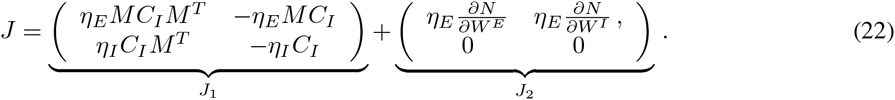

where 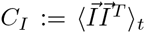 denotes the matrix of the second moments of the inhibitory inputs 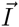 and 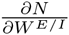 denotes the two matrices that contain the partial derivatives of the constraint term *N_i_* with respect to the excitatory and inhibitory weights 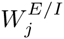, respectively. The first term *J*_1_ arises from the learning rules, the second term *J*_2_ from the normalization. For mathematical simplicity, we assume in the following that *M* is the identity matrix, i.e., that the excitatory and the inhibitory inputs are identical, although we suspect that a generalization of the derivation is straightforward. Moreover, we assume that the inputs are symmetrical in the sense that the normalised uniform vector 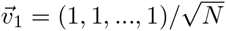 is an eigenvector of the input covariance matrix *C* and that the mean firing rates 〈*E_i_*〉 of all inputs are identical. Let *O* denote the orthogonal matrix with the eigenvectors of *C* and Λ the diagonal matrix with the eigenvalues, respectively: *C* = *O*Λ*O^T^*.

Under these assumptions, the linearized dynamics 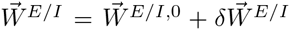 around the fixed point 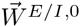 decouple almost completely when the small perturbations 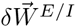 are written as a linear combination of the eigenvectors of *C*:

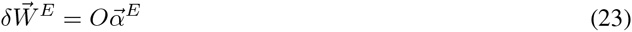

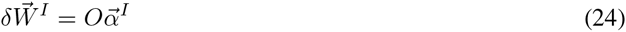

The resulting dynamical equations for the coefficient vectors 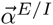 are given by

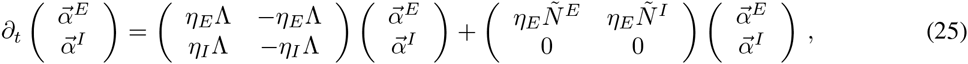

where the matrices 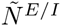 are given by 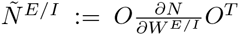. These matrices have a special structure for subtractive and multiplicative normalization. For subtractive normalization, the derivatives of the normalization term with respect to the weights is given by

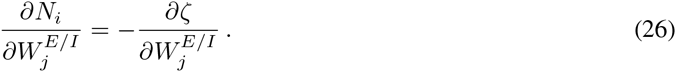

Because this term is independent of i, the product of this matrix with any vector can only generate vectors that have equal entries in all components, i.e., vectors that are proportional to 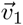. Therefore, the matrices 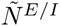 can only have non-vanishing entries in their first row.

For multiplicative normalization, the derivative of the normalization term with the respect to the weights is given by

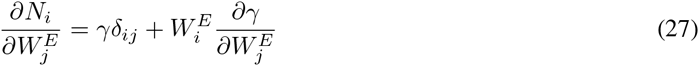

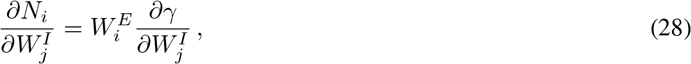

which needs to be evaluated at the fixed point 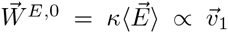 (Equation 21). If we assume that the normalization is symmetric with respect to the components of 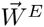, the derivative 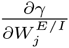 is independent of *j* at the homogeneous fixed point. As a consequence, the derivative matrices 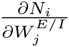 have the same eigenvectors as *C* and can therefore be diagonalized in the same basis:

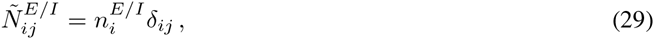

with

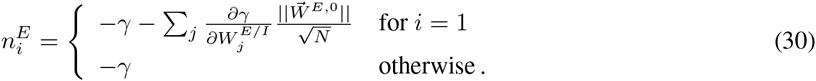

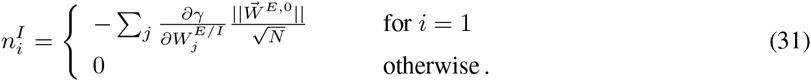

#### Stability analysis

Given the Jacobian of the learning dynamics, we can now evaluate its eigenvalues and study the stability of the fixed points. The main advantage of changing into the eigenbasis of the covariance matrix *C_I_* is that the Jacobian is almost diagonal in the sense that the only components of the excitatory and the inhibitory weights that couple belong to the same eigenvector. The only component that would have to be treated separately is the homogeneous component 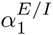. We neglect this component, however, because it is of limited interest in the context of symmetry breaking.

##### Subtractive normalization

We study the dynamics of the inhomogeneous components 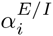 with *i* > 1, which couple only to the corresponding excitatory and inhibitory counterpart:

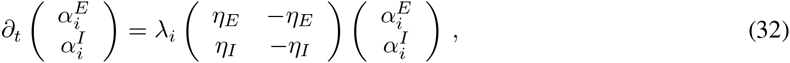

where λ_*i*_ denote the eigenvalues of the input covariance matrix *C*. The eigenvalues of this system are given by 0 (for the “balanced” eigenvector (1, 1)) and the difference between the learning rates λ_*i*_(*η_E_* − *η_I_*) (for an unbalanced eigenvector). The vanishing eigenvalue is not surprising given that the whole constraint manifold is a solution. The other eigenvalue suggests that whether a balance of excitation and inhibition is reached in finite time depends on the relation of the excitatory and inhibitory learning rates. For faster inhibitory learning, any unbalance will die out and give way to diffusive dynamics. For faster excitatory learning, all points on the constraint manifold are unstable, so that any small disruption of the E/I balance in the weights will diverge. This is only stopped by the fact that weights cannot become negative, so that the dynamics should spend most of its time in states where one excitatory weight is saturated. This state can lose stability again, however, when the inhibition had time to rebalance the excitatory weights. This is confirmed by the simulations.

##### Multiplicative normalization

The dynamics of the inhomogeneous components in the case of the multiplicative normalization, again for *i* > 1, are given by

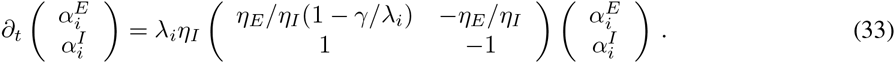

The parameters that control the stability of the fixed point are the dimensionless (and positive) ratios *g_i_* := *γ*/λ_i_ and 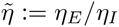. The eigenvalues 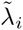 of the system can be written as a function of *g_i_* and 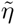:

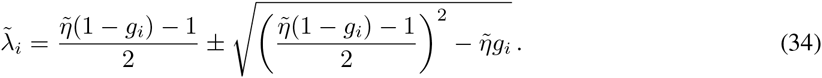

For small 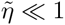, i.e., for small excitatory learning rates, the homogeneous fixed point is therefore stable. As the excitatory learning rate is increased, the eigenvalues become imaginary, until the fixed point loses stability via a Hopf bifurcation at 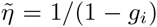. This analysis is in line with the simulations, which show the emergence of an oscillation with increasing excitatory learning rate.

